# Genetic surveillance of *Plasmodium-Anopheles* compatibility markers during *Anopheles stephensi* associated malaria outbreak

**DOI:** 10.1101/2025.03.26.645571

**Authors:** Elizabeth Waymire, Dejene Getachew, Isuru Gunarathna, Joseph Spear, Grace Lloyd, Madison Follis, Avery A. Kaye, Said Ali, Solomon Yared, Tamar E. Carter

## Abstract

Despite previous decline of malaria in Ethiopia, an outbreak in Dire Dawa in 2022 implicated invasive vector *An. stephensi* as responsible. The transmission of *Plasmodium* by invasive *An. stephensi* raises questions about the molecular basis of compatibility, and the origin of the *Plasmodium* being transmitted. The *Plasmodium P47* gene is a parasite-vector interaction gene in *Anopheles*, and along with corresponding mosquito *P47* receptor (*P47Rec*), can be critical in establishment of *Plasmodium* infections in anophelines. Here, we analyzed *P47* and *P47Rec* sequences to determine the origin of *Plasmodium* detected in *An. stephensi* during the outbreak and evaluate markers of compatibility. Analysis of geographically informative SNPs in *Pfs47* revealed that these *P. falciparum* exhibit the African haplotype. We also identified a single amino acid change in P47Rec within these *An. stephensi,* which could act as a marker for the propensity of *An. stephensi* populations to outbreaks. Together, we provide the basis for further study to deepen the understanding of invasive *An. stephensi* –African *Plasmodium* interactions to better control transmission of malaria and prevent further outbreaks.

## Introduction

*Anopheles stephensi* was first identified in Kebridehar in southeastern Ethiopia in 2016 and was detected across eastern Ethiopia soon after in 2018 [1, 2]. This vector thrives in urban environments, where it tends to lay eggs in small water pools on barrels, discarded tires, and plastic sheets around homes and construction sites [2]. The population of Addis Ababa, the capital of Ethiopia, grew from 2.377 million to 2.919 million between 2000 and 2010, later growing at a faster rate from 2.919 million to 4.705 million between 2010, and 2025 [3]. Rapid growth occurred across the country between 2010 and now, with multiple cities such as Jijiga and Dire Dawa experiencing urbanization [4, 5]. This rapid urbanization creates an ideal environment for *An. stephensi* to lay eggs, allowing the invasive populations to grow [6].

Dire Dawa city is the second most populous in Ethiopia, following the capital, Addis Ababa [7]. Dire Dawa has been experiencing rapid urbanization for the last three decades, with population increasing at a rate of 4.63% annually [8]. This urbanization has provided an environment suitable for *An. stephensi,* as demonstrated by a recent malaria outbreak. Cases in Dire Dawa were first observed in November of 2021 and continued into July of 2022. This malaria outbreak consisted of over 2,400 cases, which is over 12 times the number of cases of the last outbreak in 2019. This outbreak also continued into the dry season of Ethiopia, indicating that *An. stephensi* can thrive in urban environments in unfavorable climates [9]. In this study, surveys around the homes of malaria positive patients for mosquitoes during the outbreak revealed that 97% of the mosquitoes collected were *An. stephensi*. Statistically significant correlations were also observed between *An. stephensi* presence and *P. falciparum* positivity. This implicates *An. stephensi* as the cause of this unprecedented outbreak in Dire Dawa and highlights the major threat *An. stephensi* poses to the malaria burden in Ethiopia and beyond [9, 10].

The *Plasmodium* gametocyte surface antigen 47 (*P47)* is a gene present in female *P. falciparum* and *P. vivax.* It encodes for the P47 ligand on the surface of *Plasmodium* parasites that is expressed during the gametic and ookinete stage in the mid-gut of the mosquito [11–13]. This protein is critical for parasite-vector interactions allowing the parasite to survive the mosquito immune system and continue its infection cycle. P47 does this by suppressing the c-Jun N-terminal kinase (JNK) signaling pathway, inhibiting the production of the heme peroxidase 2 (HPX2) and NADPH-oxidase 5 (NOX5) proteins that are vital for identification of the parasites by the mosquito complement-like system [11].

Previous studies have revealed that the genetic diversity of the *P. falciparum P47* gene (*Pfs47*) is structured geographically on a continental level due to the close co-evolution between *Pfs47* and the mosquito receptor (P47Rec) of divergent anophelines. This molecular interaction has been described as a “lock-and-key” relationship [14]. Two single nucleotide polymorphisms (SNPs) have been identified at base pair locations 707 and 725 of the gene that are conserved 98-99 percent within Africa, Asia, and South America. These SNPs define three distinct haplotypes occurring between the three major continents, causing nonsynonymous amino acid (aa) mutations. These SNPs occur in Domain 2 of Pfs47, which is the most polymorphic region of the protein. Amino acid changes in this region mediate compatibility between select parasite strains and vector species [14, 15]. A study suggested that these SNPs can be used to assess the geographic origin of the *P. falciparum* and examine the risk of local transmission of imported *P. falciparum* [14]. This is particularly useful when tracking the origin of *Plasmodium* circulating during an epidemic like in Dire Dawa linked to an invasive mosquito, and estimating the propensity for further spread of the *Plasmodium.* The need to characterize the *Plasmodium* population in Dire Dawa increases, given previous evidence that this population is a hub for connectivity to other *An. stephensi* populations in eastern Ethiopia [16].

The lock-and-key relationship in the Pfs47-P47Rec system means that *P. falciparum* isolates have different compatibility to different *Anopheles* species. The highest compatibility exists between parasite and vector from the same continent, where Pfs47 acts like a key to unlock the mosquito’s immune system through the binding to P47Rec [17]. This work demonstrates that African *P. falciparum* most efficiently infects an African vector, Asian *Plasmodium* most efficiently infects an Asian vector, and so on [17]. However, a study of four single amino acid differences in African and South American *Pfs47* in the *An. gambiae* R strain and the *An. stephensi* SDA500 strain revealed that single amino acid substitutions in Pfs47 can determine mosquito infection in *An. gambiae* [18]. In *An. stephensi, P. falciparum* infection occurred regardless of the amino acid substitutions, even without Pfs47 entirely, indicating that the *Pfs47*-*P47Rec* interaction is less specific in this *An. stephensi* strain than other *Anopheles* species [18]. The strain of *An. stephensi* used in the study was the selectively bred SDA500 strain, bred to be highly susceptible to *P. falciparum* [19]. Because of this evidence, it is possible that some vectors may be naturally more permissive to many Pfs47 haplotypes [11].

Subsequent studies identified the receptor in *An. gambiae* G3 strain that interacts with Pfs47 (AgP47Rec) and found it is expressed in the mosquito midgut. This receptor has clear orthologs in other mosquito species, and sequence divergence seemed to follow speciation within anophelines [15]. This suggests that P47Rec has an ancestral basic function in anophelines and culicines, which also corroborates the continental differentiation of *Pfs47*, where *Pfs47* and *P47Rec* have been co-evolving for a long time [15]. A study showed that the highest amount of recombinant Pfs47 binding occurred between parasite and vector of shared continental origin. However, there seemed to be high levels of infectivity in South American mosquito *An. albimanus* despite low levels of Pfs47 binding, indicating there are likely other genes important for infection at play in this vector [15].

*Plasmodium vivax* evasion of the *Anopheles* immune system and parasite-vector compatibility mediation by the *P. vivax P47* gene (*Pv47*) is less understood. There are limited amino acid substitutions of the cysteine residues in Pv47, indicating a possible target for transmission blocking vaccines (TBVs). A study confirmed that Pv47 is a TBV candidate, demonstrated by transmission-reducing activity in mice using *An. dirus* [20, 21]. While confirming Pv47 as a TBV candidate, the authors also examined Pv47 sequences from El Salvador, Colombia, Vanuatu, Indonesia, Korea, India, and Thailand, discovering some indication of sequence differentiation between South America and Asia [21]. Subsequent genome-wide population analysis of *Pv47* demonstrated a clear differentiation between the eastern hemisphere and the western hemisphere parasite populations. South American *P. vivax* populations seemed to share a common ancestor and are genetically distinct from Asia. Only two African countries were represented in this study, Madagascar and Mauritania, which clustered with Indian *Pv47.* While the distribution of *P. vivax* is restricted to certain regions in Africa, limited sampling did not allow for any additional conclusions to be made about further differentiation between Africa and Asia [22].

*Plasmodium-Anopheles P47-P47Rec* mediated interaction has mostly been investigated in laboratory populations, while the nature of interaction in wild mosquitoes, especially in the context of a non-native, invasive vector, is unclear. Elucidating P47 and P47Rec sequence diversity in invasive *An. stephensi* would 1) provide vital information regarding the possibility of utilizing Pfs47 TBVs in these populations 2) reveal importation of clinically relevant *P. falciparum* from South Asia, and 3) uncover potential markers that could function as a predictor for propensity to outbreaks. Therefore, we investigated the sequence diversity of these markers in wild *An. stephensi* collected during the 2022 outbreak in Dire Dawa, providing evidence of the origin of the *Plasmodium* strains circulated by *An. stephensi* during the outbreak. We also evaluated the amino acid sequence to provide insight into the nature of *P. falciparum*-invasive *An. stephensi* compatibility. Additionally, we evaluated the nucleotide and amino acid sequences of *Pv47* in these *An. stephensi* to investigate if there are any SNPs or amino acids that separate continental origin. Finally, we sequenced the P47Rec in wild *An. stephensi* to probe intra-species diversity and gain further insight into the *Plasmodium-An. stephensi* relationship.

## Results

### Overall *Plasmodium* detection in wild *An. stephensi* from an outbreak

*P. falciparum* and *P. vivax* DNA were detected in 10.625% (17/160) of samples screened, which is a relatively high rate of detection of *Plasmodium* DNA in mosquitoes consistent with an outbreak scenario. More samples were positive for *Pfs47* than *Pv47,* and more whole-body extractions were positive than head/thorax or abdomen extractions. While whole body extractions allow more mosquitoes to be screened overall, the possibility of identifying stage of infection is lost.

### *Pfs47* diversity and relationship with African and Asian sequences

Ten samples were positive for *Pfs47*: two from abdomen extractions, two from head and thorax extractions, and six from whole body extractions (6.25% positive). Analysis of the 531 bp sequence revealed nine segregating sites contributing to eight haplotypes, with one haplotype predominately present (50%). Some chromatograms showed two nucleotides read in the same position suggesting either heterozygosity at that position or infections with multiple genetically distinct *P. falciparum* strains. Two whole body extractions were positive for both *Pfs47* and *Pv47*.

We genotyped geographically informative SNPs at positions 707 and 725 to determine continental origin of the *P. falciparum* detected in *An. stephensi.* All sequences were identified to have a C-C haplotype at SNPs 707 (236T) and 725 (242S), which is indicative of the African haplotype (Table 1). After phasing out heterozygous samples, there were 14 individual *Pfs47* sequences. In total, nine different amino acid changes were detected from the PF3D7_1346800 reference: F128L (n=1, nucleotide 382), L172S (n=1, nucleotide 515), P194H (n=14, nucleotide 581), S212R (n=1, nucleotide 636), D213G (n=2, nucleotide 638), G228A (n=1, nucleotide 683), G228V (n=1, nucleotide 683), K220E (n=1, nucleotide 658), and L248I (n=2, nucleotide 742). Specific combinations of these amino acid changes can be seen in Table 2. The sample containing the S212R mutation had no other amino acid substitutions from the reference. The most common genotype contained the P194H mutation and the I248L mutation. All the Ethiopian samples, including the positive control, had an amino acid substitution from the reference sequence (PF3D7_1346800) at the 194^th^ position, changing from P to an H. Overall, 42 segregating sites were present in the whole protein (Additional File 2: Table 2). No amino acids substitutions were observed involving cysteine residues.

**Table 1:** Alignment of *Pfs47* SNPs representative of each continent with *Pfs47* SNPs present in samples from Dire Dawa. All Dire Dawan *Pfs47* samples contain a C-C haplotype at SNP 707 and 725 indicating origin from Africa. Bolded letters denote that all samples in this study share the haplotype with Africa.

| Sample | Position |  |
| --- | --- | --- |
|  | 707 | 725 |
| *Pfs47* Papua New Guinea | T | C<sup>165</sup> |
| *Pfs47* South America | T | T |
| *Pfs47* Asia | T | C<sup>166</sup> |
| *Pfs47* Africa | C | C |
| **S220077 Dire Dawa, Ethiopia** | C | C |
| **S220098 Dire Dawa, Ethiopia** | C | C<sup>167</sup> |
| **S220108 Dire Dawa, Ethiopia** | C | C |
| **S220123 Dire Dawa, Ethiopia** | C | C<sup>168</sup> |
| **S220223 Dire Dawa, Ethiopia** | C | C |
| **S220230 Dire Dawa, Ethiopia** | C | C |
| **S220246 Dire Dawa, Ethiopia** | C | C<sup>169</sup> |
| **S220247 Dire Dawa, Ethiopia** | C | C |
| **S220267 Dire Dawa, Ethiopia** | C | C<sup>170</sup> |
| **S220282 Dire Dawa, Ethiopia** | C | C |

**Table 2:** Amino acid substitutions in Pfs47 sequences from Dire Dawa in reference to the Pf3D7 Reference Genome (Gene ID: 814213). There are eight amino acid haplotypes, as well as nucleotide haplotypes present in Dire Dawa. The number one or two after a sample ID indicates the presence of heterozygosity at that nucleotide position, resulting in two haplotypes after phasing. Amino acid substitution P194H was present in all samples, I248L in over half (n=9), and D214G in three samples. F128L, L172S, S212R, G228A, G228V, and K220E all occurred once.

| Sample ID | Amino Acid Substitutions |  |  |  |  |  |  |  |
| --- | --- | --- | --- | --- | --- | --- | --- | --- |
| S220123_1 |  |  | P194H | S212R |  |  |  |  |
| S220123_2 |  |  | P194H |  |  |  |  |  |
| S220282_1 |  |  | P194H |  | D213G | G228A |  |  |
| S220282_2 |  |  | P194H |  | D213G | G228V |  |  |
| S220267_2 | F128L | L172S | P194H |  |  |  |  | I248L |
| S220247 |  |  | P194H |  | D213G |  |  |  |
| S220246_2 |  |  | P194H |  |  |  | K220E | I248L |
| S220246_1 |  |  | P194H |  |  |  |  | I248L |
| S220223 |  |  | P194H |  |  |  |  | I248L |
| S220267_1 |  |  | P194H |  |  |  |  | I248L |
| S220230 |  |  | P194H |  |  |  |  | I248L |
| S220108 |  |  | P194H |  |  |  |  | I248L |
| S220098 |  |  | P194H |  |  |  |  | I248L |
| S220078 |  |  | P194H |  |  |  |  | I248L |

Minimum spanning network (MSN) analysis of the 312 bp sequence shows an extensive network of diversity of the Domain 2 region of the Pfs47 in Africa (Figure 1). Here, haplotypes are defined as amino acid sequences that have at least one difference. We found that Hap11 and Hap12 had the highest frequency. While there were continentally differentiated haplotypes, *Pfs47* haplotypes were shared across countries in Africa (eg. Hap11, Hap12). Dire Dawa exhibits six individual haplotypes not observed elsewhere (Hap4, Hap16, Hap17, Hap18, Hap25). Ghana shares a haplotype with Asian sequences (Hap8), and Guinea Bissau shares a haplotype with South American sequences (Hap14). Hap13 bridges the South American and Asian haplotypes (Hap14, Hap8) as well as a major African haplotype (Hap12). Hap 13 also contains samples from India, Sudan, and Papua New Guinea. The major Asian haplotype (Hap8) has two nucleotide differences compared to a haplotype present in Tanzanian and Malawi samples (Hap10), also in East Africa. South American *Pfs47* (Hap14) are conserved and distinct from other continents, and sequences from Oceania share haplotypes with all three continents (Hap8, Hap13, Hap14) (Figure 1).

**Figure 1:**
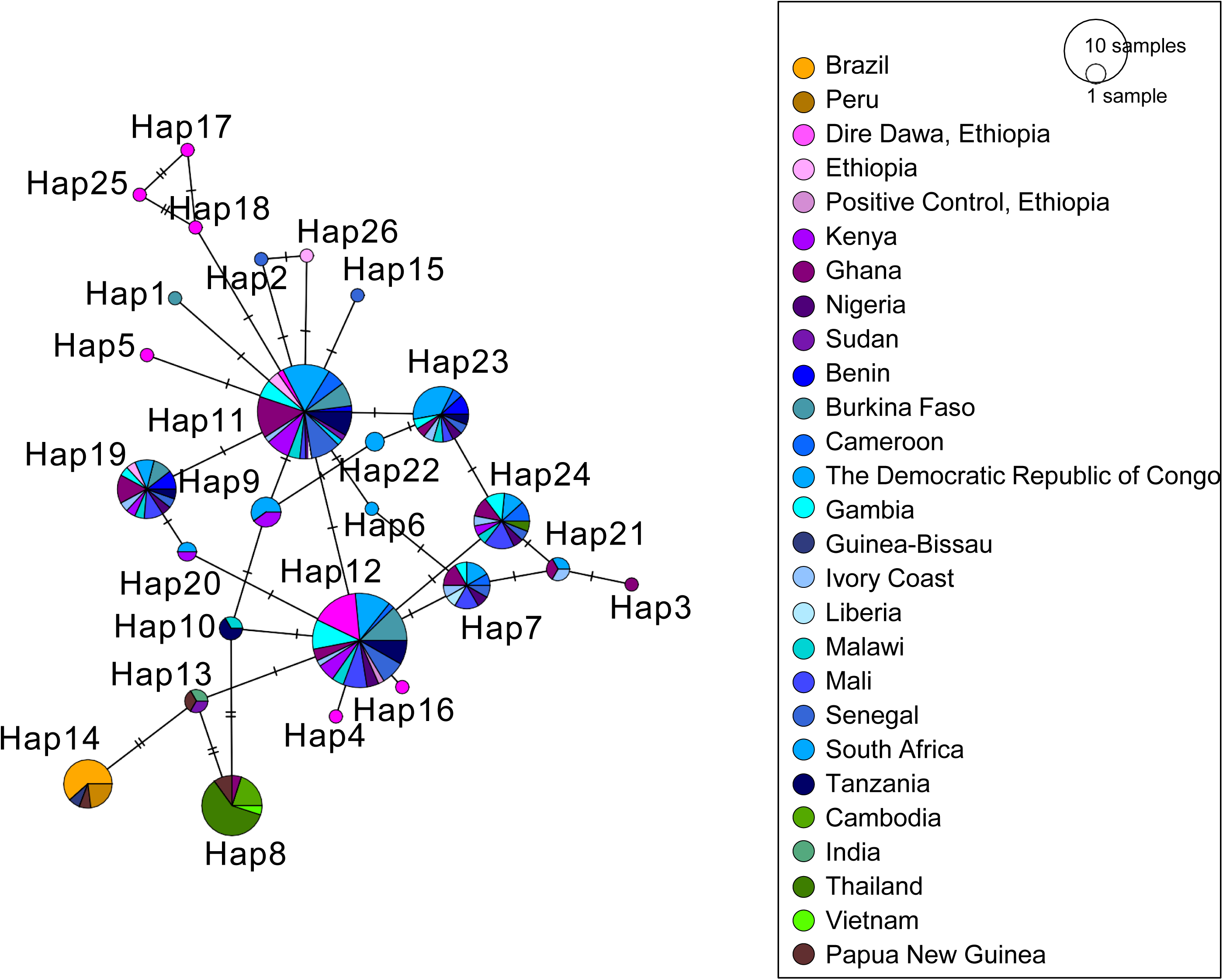
Minimum spanning network of *Pfs47* sequences from NCBI Genbank, MalariaGen database, and Dire Dawa. Samples produced in this study are represented in bright pink. Countries where invasive *An. stephensi* has been detected are represented in purple, all other countries in Africa are represented in blue, countries in Asia are represented in green, countries in South America are represented in orange, and Papua New Guinea is represented in brown. Dire Dawa shares a major haplotype with other countries in Africa (Hap11, Hap12) and has some unique haplotypes as well (Hap25, Hap17, Hap18, Hap5). Hap13 is connected to a major African haplotype (Hap12) and the South American and Asian haplotypes (Hap14, Hap8 respectively), and notably contains sequences from Sudan, India, and Papua New Guinea. Ghana shares a haplotype with the major Asian haplotype (Hap8). Thailand shares a haplotype with African sequences (Hap24).

### *Pv47* alignment

The alignment of available *Pv47* sequences from NCBI GenBank revealed three SNPs that differentiated South America from Asia and Africa at positions 1221, 1222, and 1230 of sequence JQ435617 from India 100% of the time (Additional File 1). There were no SNPs identified that differentiated all continents.

### *Pv47* diversity and relationship with African and Asian sequences

Seven samples were positive for *Pv47*: three from abdomen extractions and four from whole body extractions (2.5% positive). Four haplotypes were present, with one predominant haplotype present in four of seven samples. Two *Pv47* positive samples indicated either heterozygosity or multiple different *P. vivax* strains.

There is a single haplotype that contains most of the samples from Africa and Asia (Hap3). Dire Dawa has another haplotype that is differentiated from other Africa haplotypes (Hap11). South American haplotypes (Hap5 and Hap6) are the most like a haplotype in Thailand (Hap2) but with several nucleotide differences separating them. Sequences from Oceania share the major African and Asian haplotype (Hap3) and have one unique haplotype (Hap10) (Figure 2). A second major Asian haplotype (Hap8) comprised of mostly India, Pakistan, and Afghanistan shares sequence similarity with one sequence from Ethiopia (Figure 2). All the Pv47 samples had amino acids changes from the reference sequence (XM001614197.1): F22L, F24L and K27E. These amino acid substitutions have been observed previously in Asian sequences, although there are 21 segregating sites in the whole protein globally (Additional File 2: Table 3) [21]. No amino acids substitutions were observed involving cysteine residues.

**Figure 2:**
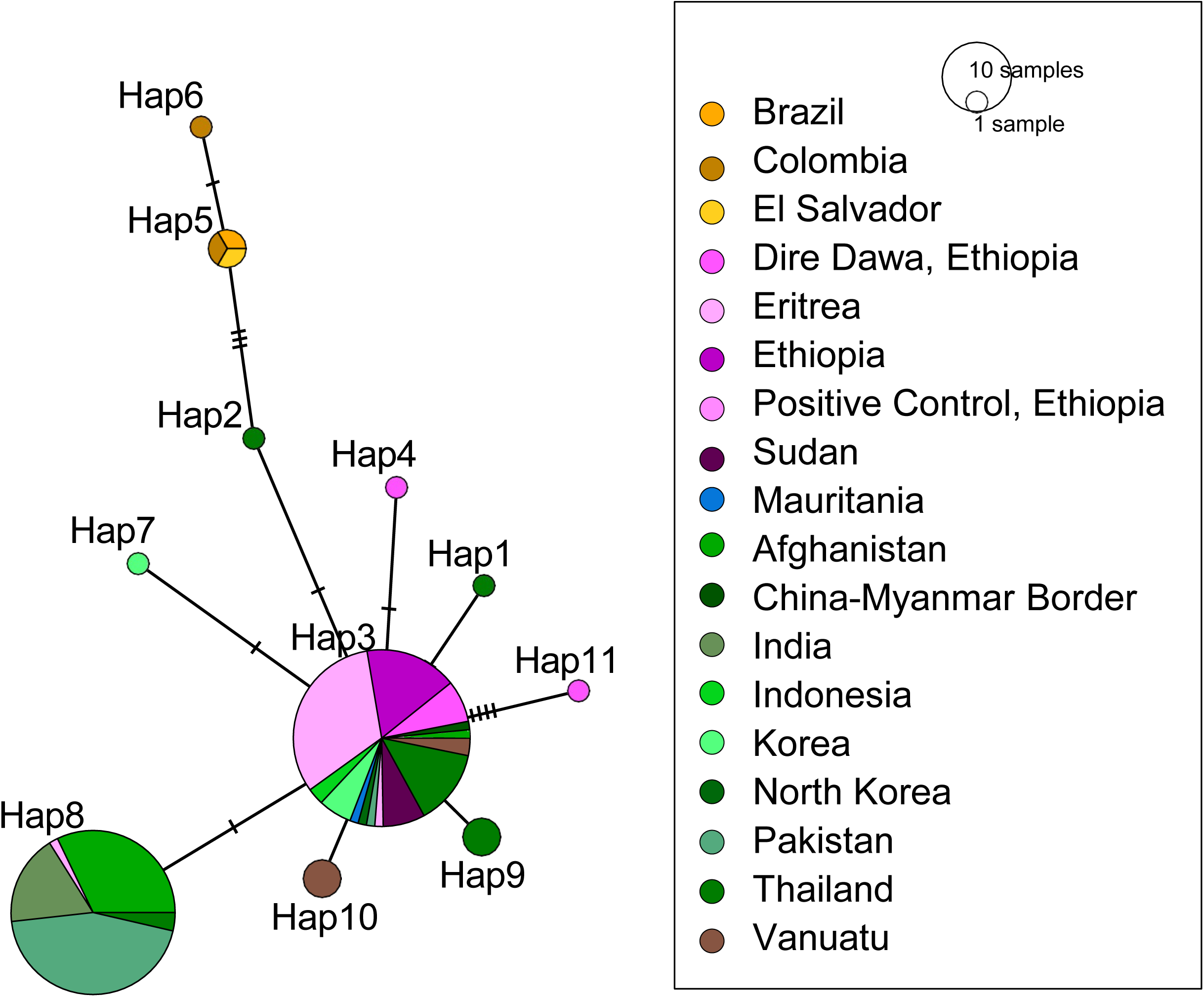
Minimum spanning network of *Pv47* sequences from NCBI and Dire Dawa. Samples produced in this study are represented in bright pink. Countries where invasive *An. stephensi* has been detected are represented in purple, all other countries in Africa are represented in blue, countries in Asia are represented in green, countries in South America are represented in orange, and Vanuatu is represented in brown. Hap3 represents most sequences from Africa and Asia, although Hap8 is also a major Asian haplotype also shared with the positive control from Ethiopia. South American sequences are present in Hap6 and Hap7. Samples from Dire Dawa share similarity with Hap3 but have two unique haplotypes (Hap11, Hap4).

**Table 3:**
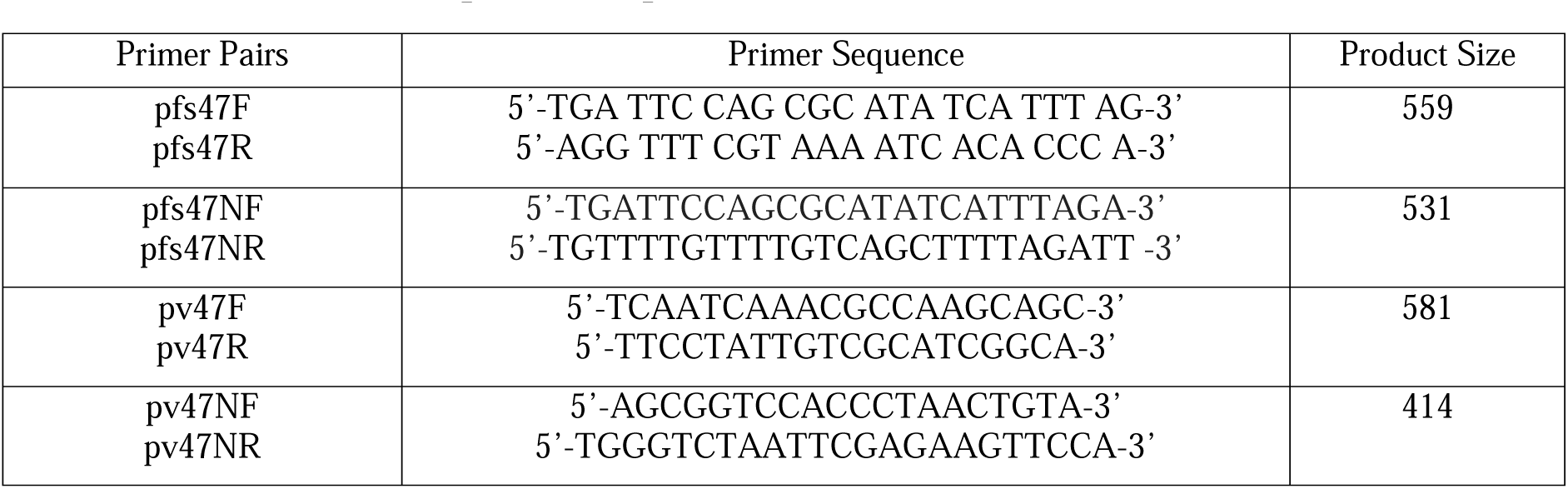
*Pfs47* and *Pv47* primer sequences.

### P47Rec diversity

We sequenced the entire *P47Rec* gene in a subset of samples to evaluate *An. stephensi* diversity in multiple populations across time that have demonstrated increased malaria transmission or significance in gene flow within the Horn of Africa [16, 23]. The P47Rec amino acid sequences were the same in all *An. stephensi* from Ethiopia except for one amino acid in the second exon (aa position 53), and the second exon was sequenced for the rest of the samples. At this position, samples were either homozygous for histidine (53H) or glutamine (53Q) or were heterozygous, consistent with *An. stephensi* reference sequences. In 2018, Kebridehar had two heterozygous samples (2/16) and the rest were homozygous for H (14/16), Semera had three heterozygous samples (3/17) and the rest were homozygous for H (14/17), and Dire Dawa had two samples homozygous for Q (2/23), seven heterozygous samples (7/23), and the rest homozygous for H (14/23). In Lawyacado, one sample was homozygous for Q, two were heterozygous (2/11) and the rest were homozygous for H (8/11). In Dire Dawa in 2022, all *P. falciparum* positive samples from this study were homozygous for H (6/6) with samples positive for both *P. falciparum* and *P. vivax* being both homozygous for H (1/2) and heterozygous (1/2). The non-positive samples from Dire Dawa 2022 had one sample homozygous for Q (1/19), five heterozygous samples (5/19) and the rest were homozygous for H (13/19). Samples from India had homozygosity for both H and Q (12/20 HH, 1/20 QQ) and seven heterozygous samples (7/20) (Figure 3, Additional File 2: Table 4).

**Figure 3:**
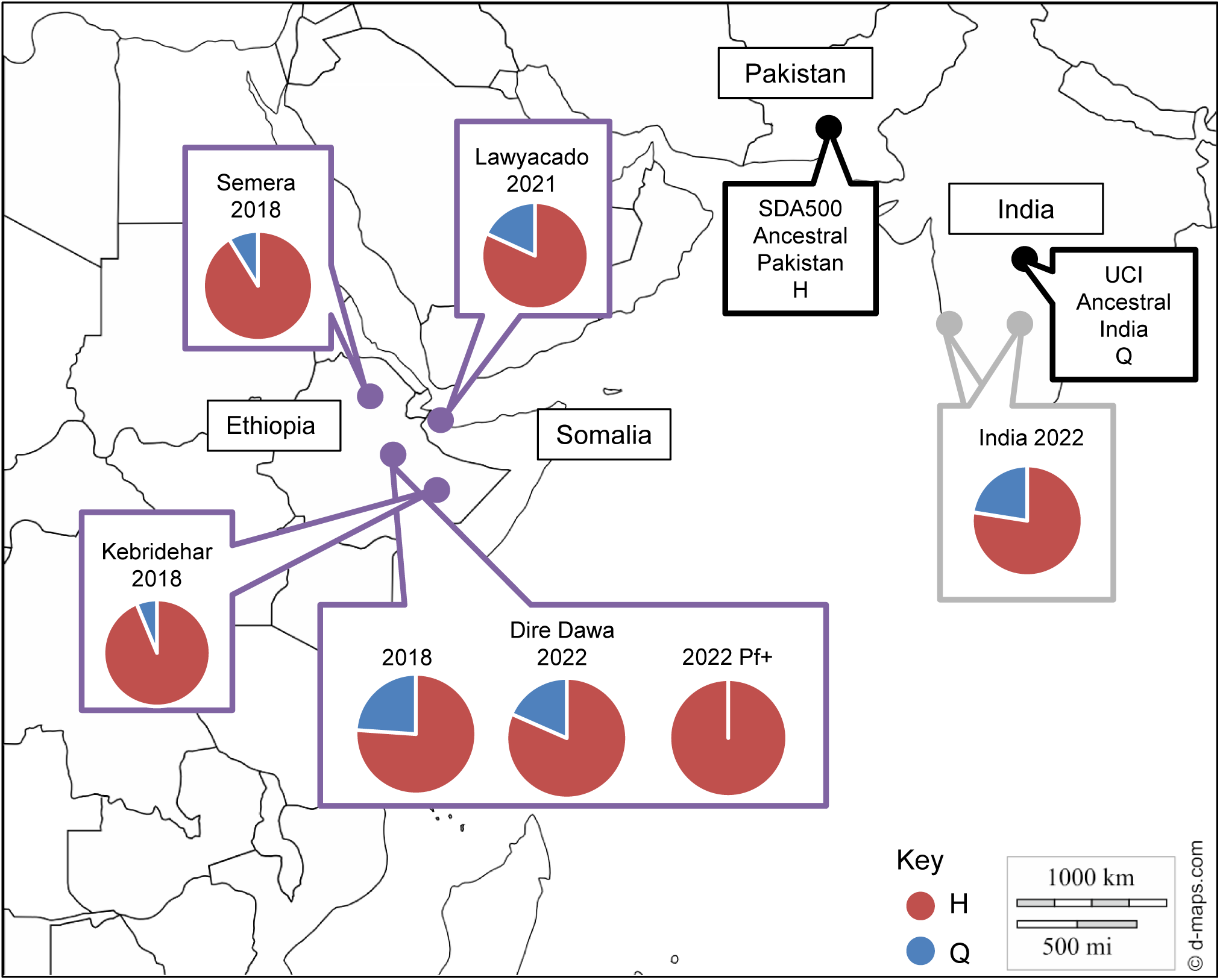
Allele frequency of amino acids histidine or glutamine at position 53 in different samples and reference genomes of *An. stephensi*. Invasive populations of *An. stephensi* are outlined in purple, native in gray, and reference in black. Allele frequency of H is denoted by red in the pie charts and frequency of Q is denoted by blue.

### *Pfs47/Pv47* and *COI* diversity

*COI* haplotypes were evaluated in the samples to investigate any *An. stephensi* lineages specifically infected by *P. falciparum* or *P. vivax*. Four previously identified *COI* haplotypes from Ethiopia were present in the sample set. Seventy-four samples were identified as Haplotype 2, ten as Haplotype 4, one as Haplotype 1, and 23 as Haplotype 3. Haplotype 2 samples had the most positive samples but were also the most prevalent, Haplotypes 2, 3, and 4 were observed in malaria positive samples, with the most positives being observed in Haplotype 2. Haplotype 2 was the most predominant haplotype, so a larger sample size is needed to determine any correlation (Additional File 2: Figure 1).

## Discussion

The data presented here provides novel evidence that the *P. falciparum* present in the invasive *An. stephensi* collected during an outbreak were of African origin. In *Pfs47*, the SNPs at locations 707 and 725 of the gene revealed that the *P. falciparum* in these *An. stephensi* exhibit the genotype that is conserved in 99.8% of African *Pfs47* [14]. All *Pfs47* haplotypes identified in this study were either connected with or shared haplotypes with other African sequences, further confirming African origin of the *P. falciparum* in these *An. stephensi*. Additional support relates to previously identified markers for artemisinin and diagnostic resistance in the *P. falciparum* during the outbreak, where two mutations were identified that have previously been found in the Horn of Africa [9]. Together, these data provide evidence to support that the *P. falciparum* transmitted by *An. stephensi* during the outbreak was of African origin, rather than importing South Asian *P. falciparum* and emphasizes a variation among anophelines and their specificity.

While most of the African *P. falciparum* haplotypes clustered separately from other continental groups, the MSN analysis for *Pfs47* revealed a connection between Asian and African haplotypes. The predominant Asian *Pfs47* haplotype (Hap8) has the closest connection to Hap 10, which includes Tanzania and Malawi, and Hap13, which includes Sudan and India. Population genetics analysis of Sudanese *An. stephensi* have shown considerably high genetic variation consistent with being one of the older invasive *An. stephensi* populations [24]. Finding shared *Pfs47* haplotypes with India highlights the connection between Sudan and South Asia, and the role the location may have played in the earlier introduction of *An. stephensi*. Further studies on the *P. falciparum* diversity across the *An. stephensi* range could provide additional insight into the potential for importation.

In other vectors, like local *An. gambiae* sensu stricto (s.s), *P. falciparum* infection is based on a lock-and-key theory where the parasite has a variable region in *Pfs47* that allows for evasion of the mosquitos’ immune system [13, 17]. In these wild *An. stephensi,* eight *Pfs47* haplotypes were observed with some samples containing the I248L mutation previously identified. A study found that in *An. gambiae sensu stricto,* this mutation caused 100% melanization of parasites, but in the *An. stephensi* SDA500 there were 100% live parasites indicating that singular amino acid changes, whether conservative or not, do not impact infection. Presumably, with both the isoleucine and leucine present in this *An. stephensi* parasite population, it seems as though singular amino acid mutations may determine infection in local African vectors but not define infection in invasive *An. stephensi*. Nucleotide diversity in the *Pfs47* gene across human infections has been observed previously within a single country of origin. The presence of multiple amino acid haplotypes within a single vector in one city suggests a more ambiguous interaction between P47 and P47Rec than in *An. gambiae* complex mosquitoes [14, 18]. Significantly, no amino substitutions were observed involving cysteine residues in either Pfs47 or Pv47, indicating the continued possibility for these genes to be an effective TBV target in this region.

The P47Rec of other anopheline vectors with well-studied Pfs47*-*P47Rec relationships, such as *An. gambiae,* are 100% conserved across the African continent [25]. This supports the lock-and-key theory, where *An. gambiae-P. falciparum* interaction is very specific. We see in these invasive *An. stephensi* that there is genetic diversity across time and across the Horn of Africa, supporting the idea that *An. stephensi* has more ambiguous *Pfs47* interaction with *P. falciparum* [11]. Moreover, all genotypes present in the *P. falciparum* positive *An. stephensi* were homozygous for histidine. When both *P. falciparum* and *P. vivax* were present, glutamine was only present in heterozygous form, potentially indicating that population level diversity in *An. stephensi* may influence variation within *Pfs47*. A larger sample size accompanied by experimental infection studies is needed to make definitive conclusions.

There is limited data available on *P. vivax Pv47* related to basic biology, compatibility with different vectors, and geographical specificity. Using data from NCBI GenBank, we were unable to identify SNPs or amino acid changes that differentiate continental origin between Africa and Asia, agreeing with a previous study on global *P. vivax* found in both African and Asian samples [22]. Dire Dawa *Pv47* exhibits more diversity than other African sequences, although there is only representation from three African countries. There is a distinct haplotype present in Dire Dawa (Hap4) that shows distant relatedness to the rest of African and Asian Hap3.

Overall, the story of *Pfs47* mediated *Plasmodium* infection in *An. stephensi* seems to be much more complex than in native African anophelines. To this end, we present compelling data on both the vector and the parasite, *P47* and *P47Rec*, in which there is more diversity present in a singular species than observed in local malaria vectors like the *An. gambiae* complex. The results of this study emphasize that invasive *An. stephensi* may not strictly follow the lock-and-key model where a high number of conservative and non-conservative Pfs47 amino acid changes are observed as well as a receptor that does not appear to be conserved, as previously noted [11]. This is indicated by the agreeance with laboratory data of *An. stephensi* (SDA500), where there are multiple amino acid changes in Pfs47 that did not affect *P. falciparum* infectivity. While we cannot confirm that the *Plasmodium* DNA detected here was not of melanized parasites, the discovery of these diverse *P. falciparum Pfs47* sequences are still alarming, considering it could be suggestive that wild *An. stephensi* is like the laboratory strain in the P47-P47Rec system. Limited information can be concluded about *Pv47.* However, it appears *Pv47* from Africa and Asia are not clearly delineated, which could be explained by shared recent ancestry [22]. If P47 is a marker of compatibility in *P. vivax* like it is in *P. falciparum*, then the similarity in Asian and African ligand would indicate ease of *P. vivax* infection by both invasive and local anophelines. This information provides evidence for the need for further study in *An. stephensi* to determine what other genes are involved in determining both *P. falciparum* and *P. vivax* infection in these invasive populations. This will aid in finding targets for future malaria control as well as continue to investigate this potential marker that could function as a predictor for propensity to outbreaks.

## Methods

### Collection of *An. stephensi*

*Anopheles stephensi* were collected in Dire Dawa, Ethiopia during March of 2022. Adult mosquitoes were caught from indoor offices or outdoor manholes via mouth aspirator. Mosquitoes were sorted between culicines and *Anopheles.* The latter were discriminated according to a morphological key [26]. *An. stephensi* mosquitoes were sexed, and females were stored individually in 1.5 mL tubes in a bag containing silica gel. It was denoted which mosquitoes had engorged abdomens and they were labeled as “blood fed.”

### DNA extraction and molecular identification of mosquitoes

All mosquitoes that were identified as recently blood fed were extracted (n=5), and the rest of the mosquitoes were selected at random (n=155). DNA was extracted from either the head and thorax (n=51), abdomen (n=51), or whole body (n=58) using the DNeasy Blood and Tissue Kit or the DNA Micro Kit (Qiagen, Valencia, CA). Detection of malaria in the head and thorax or abdomen could suggest a particular state of *Plasmodium* development, but whole body detection allows for more mosquitoes to be screened. Polymerase chain reaction (PCR) was conducted for each mosquito, targeting the *An. stephensi* specific nuclear internal transcribed spacer 2 (*ITS2*) locus and a universal mitochondrial cytochrome *c* oxidase subunit 1 (*COI*) [2, 27]. The final reagent components and concentrations for PCR were 1X Promega G2 HotStart Master Mix (Promega, Madison, WI), 0.5 mM for both primers, and 1 uL of DNA template. The endpoint assay targeting the *ITS2* locus of *An. stephensi* [2]. PCR temperature protocol was 95°C 1 min, 30 cycles of 95°C for 30s, 48°C for 30 s, and 72°C for 1 min; followed by 72°C for 10 min. The PCR product was visualized via gel electrophoresis, and a 522-base pair (bp) band was identified. Only *An. stephensi* samples would contain a band, so any samples that did not produce a band were not included in the sample set. PCR temperature protocol consisted of 95°C 1 min, 30 cycles of 95°C for 30 s, 48°C for 30 s, and 72°C for 1 min; followed by 72°C for 10 min. *COI* PCR products were sequenced using Sanger technology by commercial laboratory (Eurofins Genomics LLC, Psomagen).

### *Pv47* alignment for SNP determination

An alignment was performed using CodonCode v9.01 to determine if there were any SNPs or amino acid substitutions that determined continental differences. Sequences of *Pv47* were first downloaded from National Center for Biotechnology and Information (NCBI) GenBank, and previously generated *Pv47* sequences from Ethiopia were included. The sequences were opened in CodonCode and the sequences were aligned. Sequences were organized by continent and differences in the sequence were visualized. Sequences were then translated into amino acids and the reading frame was selected based on the reference amino acid sequence in NCBI. Differences in amino acids were then visualized.

### *Pfs47* and *Pv47 P. falciparum* and *P. vivax* primer design

For the *Pfs47* primers, the *Pfs47* gene from the reference *P. falciparum* genome (NCBI: taxid36329) was used. An alignment of available *Pfs47* sequences was performed and the SNPs identified in Molina-Cruz et al. 2021 to differentiate continental origin were identified in the alignment [14]. The reference gene was put into Primer3Plus, with a target product indicating where the SNPs were identified to be. Primers were tested using a positive control of *P. falciparum* DNA. Positive control amplicons of *Pfs47* were sequenced to confirm the correct target was amplified, amplifying a 559 bp target (Table 3).

For the *Pfs47* nested primers, the amplified sequence of the un-nested primers was determined using NCBI Primer Blast. This product was uploaded to Primer3Plus and the area previously identified with SNPs was selected as a target. Potential primers were tested against the non-redundant and RefSeq mRNA databases in NCBI Primer Blast, and primers that didn’t also target *An. stephensi* DNA were selected. These primers were tested on the positive control DNA and sequenced to confirm the correct target. This produced a product of 531 bp (Table 3).

A similar protocol was used to design the *Pv47* and *Pv47* nested primers. The *Pv47* gene from the reference *P. vivax* genome (NCBI: txid126793) was used. The gene was put into NCBI Primer Blast excluding *An. stephensi* DNA with a target indicating where SNPs were identified to be in the described alignment. Potential primers were tested against the non-redundant and RefSeq mRNA databases in NCBI Primer Blast and primers that didn’t also target *An. stephensi* DNA were selected. Primers were tested using a positive control of *P. vivax* DNA. Positive control amplicons were sequenced to confirm the correct target was amplified, amplifying a 581 bp target (Table 3).

The *Pv47* nested primers were created by first determining the amplified sequence of the un-nested primers using NCBI Primer Blast. The product was uploaded to Primer3Plus and the area previously identified to contain the SNPs was selected as a target. Potential primers were tested against the non-redundant and RefSeq mRNA databases in NCBI Primer Blast, and primers that did not also target *An. stephensi* DNA were selected. These primers were tested on the positive control and sequenced to confirm the correct target, a product of 414 bp (Table 3).

### *Pfs47* based *P. falciparum* detection

PCR was conducted for the *Pfs47* gene to detect *P. falciparum* presence in *An. stephens*i samples. A positive control of positive *P. falciparum* human blood DNA extractions (provided by Eugenia Lo at the University of North Carolina at Charlotte) served to verify successful amplification, alongside a negative control lacking genomic DNA to ensure no contamination was present. Un-nested PCR reactions were conducted initially to detect *P. falciparum Pfs47* presence before running nested PCR reactions. The *Pfs47* primers amplified a 559 bp un-nested fragment in *P. falciparum* (Table 3). Unnested protocol reagents and concentrations consisted of 1X Promega G2 HotStart Master Mix (Promega, Madison, Wisconsin, USA), 0.4 mM of primer, plus 4 uL of isolated DNA template. The cycling protocol was as follows: 95°C for 1 min, 34 cycles of 95°C for 1 min, 57°C for 1 min, 72°C for 1.5 min, and an extension step of 72°C for 10 min. The nested protocol called for a second set of primers to selectively amplify a 531 bp fragment of *P. falciparum Pfs47* (Table 3). The nested reaction was performed with 1X Promega G2 HotStart Master Mix (Promega, Madison, Wisconsin, USA) 0.4 mM of primer, plus 2 uL of the PCR product from the initial un-nested reaction. The cycling protocol was as follows: 95°C for 10 min, 34 cycles of 95°C for 1 min, 58°C for 1 min, 72°C for 1 min, followed by 72°C for 5 min.

### *Pv47* based *P. vivax* detection

PCR was conducted to amplify the *Pv47* gene for the detection of *P. vivax* in mosquito samples. A positive control of positive *P. vivax* human blood DNA extractions (provided by Eugenia Lo at the University of North Carolina at Charlotte) was included, alongside a negative control lacking genomic DNA to ensure no contamination was present. An un-nested protocol was used to detect the presence of *P. vivax Pv47,* followed by a nested protocol.

*Pv47* primers amplified a 581 bp un-nested fragment in *Plasmodium vivax* (Table 3). The un-nested protocol reagents and concentrations consisted of 1X Promega G2 HotStart Master Mix (Promega, Madison, Wisconsin, USA), 0.4 mM of primer, plus 4 uL of isolated DNA template. The cycling protocol was as follows: 95°C for 1 min, 34 cycles of 95°C for 1 min, 58°C for 1 min, 72°C for 1.5 min, and an extension step of 72°C for 10 min. The nested protocol called for a second set of primers to selectively amplify a 414 bp fragment of *P. vivax Pv47* (Table 3). The nested reaction was performed with 1X Promega G2 HotStart Master Mix (Promega, Madison, Wisconsin, USA), 0.4 mM of primer, plus 2 uL of the PCR product from the initial un-nested reaction. The cycling protocol was as follows: 95°C for 10 min, 34 cycles of 95°C for 1 min, 60°C for 1 min, 72°C for 1 min, followed by 72°C for 5 min. All PCR products were run on a 2% agarose gel and visualized. Positive samples were sequenced via Sanger Sequencing at a commercial laboratory (Eurofins Genomics LLC, Psomagen).

### *P47Rec* Amplification

PCR was conducted to amplify the P47 Receptor in *An. stephensi* samples. 20 mosquitoes were selected from Kebridehar, Ethiopia (2018), 20 from Semera, Ethiopia (2018), 20 from Dire Dawa, Ethiopia (2018), 13 from Lawyacado, Somalia (2021), and 20 more from Dire Dawa (2022). P47Rec primers amplified a 688 bp fragment including the second and third exon of the protein (F: 5’-TGGCAAATGACTAACGTGGA-3’, R: 5’-GTGTTGCCAGTTCGCTGTAA-3’). The cycling protocol was as follows: 95°C for 1 min, 34 cycles of 95°C for 1 min, 58°C for 1 min, 72°C for 1.5 min, and an extension step of 72°C for 10 min. All PCR products were run on a 2% agarose gel and visualized; all samples were sequenced via Sanger Sequencing at a commercial laboratory (Psomagen).

### *Pfs47* and *Pv47* sequence analysis

The haplotypes for the sequences that indicated multiple nucleotides read at a single position were phased out using DNAsp v 5.10.01. The .FASTA file containing cleaned sequences for both genes were input into DNAsp and phased using MCMC standard options (100 iterations, 1 thinning interval, 100 burn-in iterations). A new .FASTA file was exported and each sample that had multiple haplotypes was denoted as haplotype 1 or 2 after the sample name. Sequences were also translated using CodonCode v9.0.1 and the reading frame was selected to match the protein coding of Pfs47 and Pv47 on NCBI (PV077639-PV077662).

### *COI* phylogenetic analysis

Phylogenetic analysis was first performed on the *COI* sequences of *Pfs47* or *Pv47* positive samples with an outgroup of *An. maculatus* (KT382822). Phylogenetic analyses were estimated using a maximum likelihood approach with RaxML. The GTR GAMMA option that uses the general time-reversible model of nucleotide substitution with the gamma model rate of heterogeneity was used. 1000 runs were completed with rapid bootstrap analysis. The RAxML output was viewed in FigTree with a root at the outgroup.

### Isolation of Pfs47, Pv47, and P47Rec Sequences from Whole Genome Data

The extraction of Pfs47, Pv47, and P47Rec sequences followed a comprehensive bioinformatics pipeline. First, the raw FASTQ files were obtained from databases MalariaGen and NCBI [28, 30]. These sequences were aligned to their respective reference genomes (PF3D7 for *Pfs47,* XM_001614197 for *Pv47*, and UCI_ANSTEP_V1.0 for *P47Rec*) using Bowtie2, ensuring precise mapping of reads. The aligned sequences were converted into BAM files using Samtools to create binary alignments. Subsequently, Bcftools was used for variant calling, employing ‘mpileup’ to aggregate aligned reads and ‘call’ to identify potential variants. Variant filtration ensured high-quality data, with filters applied for Phred-scaled quality scores greater than 30, read depths above 10, and variant frequencies above 1%.

After quality filtering, the variants were processed to generate FASTA sequences. These sequences were further aligned using MAFFT to prepare them for downstream analyses, including evolutionary or functional assessments [29]. This methodical process ensured accurate extraction and alignment of the target gene sequences from complex whole genome datasets.

### Minimum spanning network analysis

Sequences from online databases like NCBI GenBank and MalariaGen were processed as described and aligned with all *Pfs47* or *Pv47* sequences in this study using CodonCode Aligner v9.0.1 and exported into a FASTA file. This FASTA file was converted to .rphylips and the data was copied into Microsoft Excel. Random haplotype names were given to every unique haplotype for both *Pfs47* and *Pv47*. The frequencies and sequences of each haplotype were formatted into a .nex file to be imported in popart [31]. A minimum spanning network was then created using standard settings and an epsilon value of 0. Haplotypes were colored according to the continent and the presence of *An. stephensi*.

## Abbreviations

JNK: c-Jun N-terminal kinase
HPX2: heme peroxidase 2
NOX5: NADPH-oxidase 5
TBV: Transmission Blocking Vaccine
MSN: Minimum spanning network
P47Rec: P47 receptor
SNPs: single nucleotide polymorphisms
PCR: polymerase chain reaction
*ITS2*: internal transcribed spacer 2
*COI*: cytochrome *c* oxidase subunit 1
NCBI: National Center for Biotechnology Information

## Additional Files

Additional File 1.xls: *Pv47* global variant sites. This file contains a table of data with the position of each variant across publicly available data and data found here. Sites 1221, 1222, and 1230 are bolded to demonstrate that the variation at these sites separates Asia and Africa away from South American *Pv47*.

Additional File 2.doc:

Additional File Figure 1: Phylogenetic tree of *An. stephensi COI* colored by *pfs47* or *pvs47* positive status. Only *COI* haplotypes 2, 3, and 4 were present in *Plasmodium* positive samples. Blue samples represent *pfs47* positive samples, *pvs47* positive samples are represented by red color, and samples positive for both *pfs47* and *pvs47* are represented by purple color. Different shades of red and blue indicate different *pfs47* or *pvs47* haplotypes, and samples with multiple haplotypes are designated by arrows to differentiate the haplotypes present.

Additional File Table 2: Segregating sites in Pfs47 globally. This table contains data of all segregating sites present in the Pfs47 protein across publicly available data along with the allele frequency. Amino acids in bold are those found in samples in this study. ND represents no data.

Additional File Table 3: Segregating sites in Pv47 globally. This table contains data of all segregating sites present in the Pv47 protein across publicly available data along with allele frequency. Amino acids in bold are those found in samples in this study. ND represents no data. Additional File Table 4: Presence of homozygous allele and allele frequency of histidine and glutamine at amino acid position 53 in P47 receptor over time.

## Declarations

### Ethics approval and consent to participate

Not applicable.

### Consent for publication

Not applicable.

### Availability of data and materials

All data generated or analyzed during this study are included in this published article and its supplementary information files. Sequences for *Pfs47* and *Pvs47* are published in NCBI GenBank under accession numbers PV077639-PV077662, *P47Rec* sequences are under accession numbers PV100900 – PV101033 and PV179422-PV179438.

### Author contributions

EW, IG, and TEC contributed to the conceptualization of the project. EW and TEC contributed to the design of the study. SY and DG designed the mosquito collection protocol. DG and SA collected the mosquitoes. EW and IG conducted literature search. EW, IG, MF, GL, JS, and AK contributed to data production and molecular analysis. EW, IG, and TEC contributed to figure creation. EW, IG, AK, GL, and TEC contributed to the writing of the manuscript. MF, AK, GL, IG, DG, and SY contributed to manuscript revision. All authors read and approved the final manuscript.

## Acknowledgements

We acknowledge the support of Eugenia Lo for supplying the positive control for both *P. falciparum* and *P. vivax*. We also acknowledge the support of Sarah Zohdy for aiding in discussions of this project.

