## Additional File 2 for "Genetic surveillance of *Plasmodium-Anopheles* compatibility markers during *Anopheles stephensi* associated malaria outbreak"

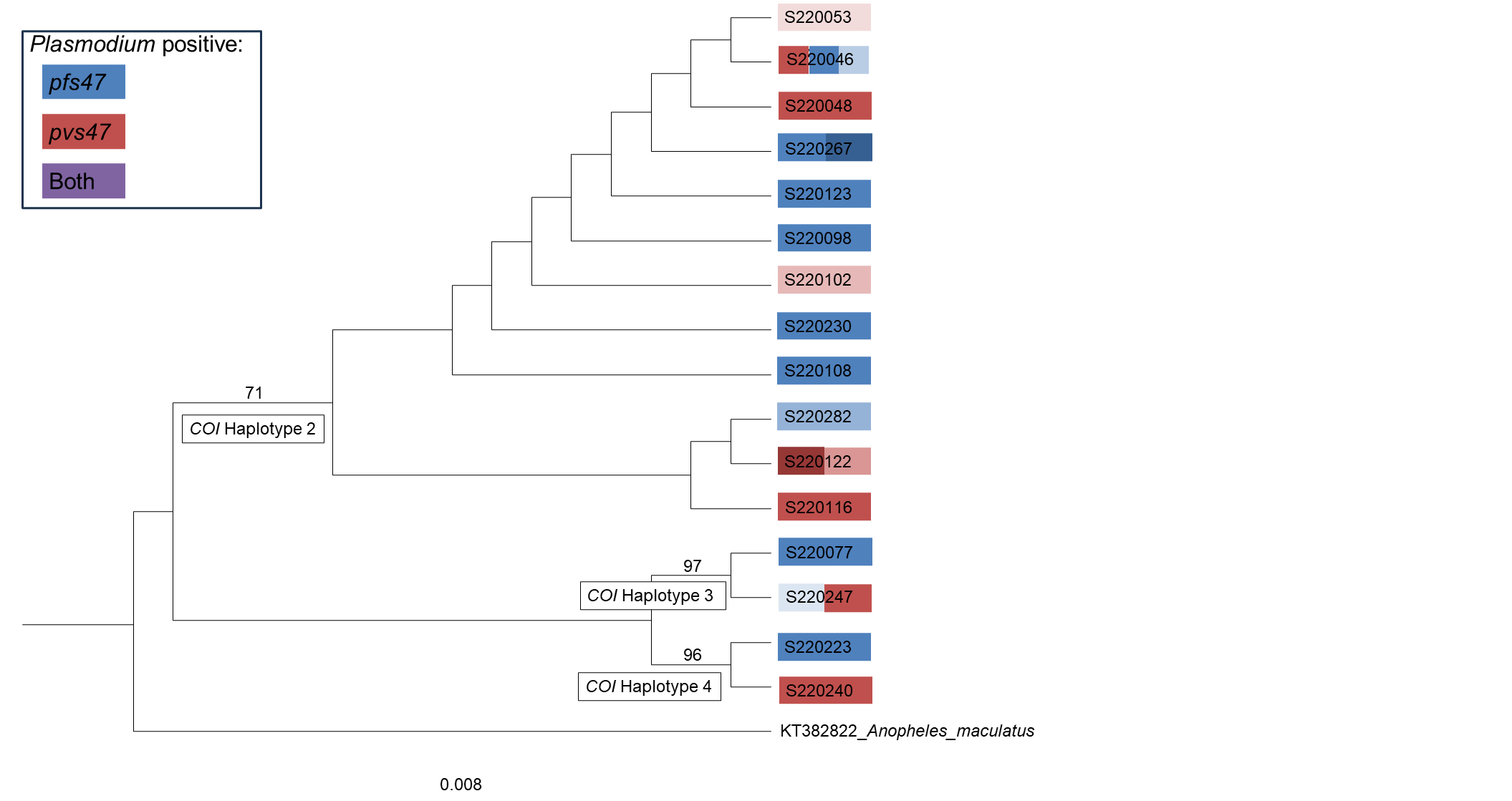

**Figure 1.** Phylogenetic tree of *An. stephensi COI* colored by *pfs47* or *pv47* positive status. Only *COI* haplotypes 2, 3, and 4 were present in *Plasmodium* positive samples. Blue samples represent *pfs47* positive samples, *pvs47* positive samples are represented by red color, and samples positive for both *pfs47* and *pv47* are represented by purple color. Different shades of red and blue indicate different *pfs47* or *pv47* haplotypes, and samples with multiple haplotypes are designated by arrows to differentiate the haplotypes present.

**Table 2:** Segregating sites in Pfs47 globally. Amino acids in bold are those found in samples in this study. ND represents no data.

| **Position** | **Amino Acid** | **Freq.** | **Amino Acid** | **Freq.** | **Amino Acid** | **Freq.** |
| --- | --- | --- | --- | --- | --- | --- |
| 27 | D | 0.08849558 | E | 0.42920354 | ND | 0.48230088 |
| 28 | L | 0.48230088 | I | 0.08849558 | ND | 0.42920354 |
| 46 | L | 0.00442478 | V | 0.7920354 | ND | 0.20353982 |
| 55 | K | 0.06637168 | E | 0.7300885 | ND | 0.20353982 |
| 106 | N | 0.87168142 | S | 0.00442478 | ND | 0.12389381 |
| 110 | T | 0.00442478 | P | 0.00442478 | ND | 0.12389381 |
| 128 | L | 0.00442478 | F | 0.93362832 | ND | 0.0619469 |
| 129 | R | 0.00442478 | S | 0.93362832 | ND | 0.0619469 |
| 144 | N | 0.00884956 | S | 0.98230088 | ND | 0.00884956 |
| 155 | N | 0.00884956 | S | 0.98230088 | ND | 0.00884956 |
| 165 | N | 0.00442478 | D | 0.99115044 | 'Y': | 0.00442478 |
| 172 | L | 0.99557522 | S | 0.00442478 |  |  |
| 188 | D | 0.17699115 | E | 0.82300885 |  |  |
| 194 | H | 0.90265487 | P | 0.09734513 |  |  |
| 212 | R | 0.00442478 | S | 0.99557522 |  |  |
| 213 | D | 0.98672566 | G | 0.01327434 |  |  |
| 219 | I | 0.07522124 | M | 0.92477876 |  |  |
| 220 | K | 0.99557522 | E | 0.00442478 |  |  |
| 224 | N | 0.08849558 | V | 0.00442478 | 'I': | 0.90707965 |
| 228 | G | 0.99115044 | V | 0.00442478 | 'A': | 0.00442478 |
| 236 | I | 0.15486726 | T | 0.84513274 |  |  |
| 240 | L | 0.86725664 | I | 0.13274336 |  |  |
| 242 | L | 0.05752212 | S | 0.94247788 |  |  |
| 247 | V | 0.94247788 | A | 0.05752212 |  |  |
| 248 | L | 0.54867257 | I | 0.45132743 |  |  |
| 262 | I | 0.00442478 | ND | 0.02212389 | N | 0.00442478 |
| 266 | D | 0.96460177 | E | 0.00884956 | ND | 0.02654867 |
| 270 | Q | 0.00442478 | E | 0.96902655 | ND | 0.02654867 |
| 272 | I | 0.21681416 | ND | 0.02654867 | N | 0.57522124 |
| 273 | N | 0.00442478 | H | 0.00442478 | Y | 0.96460177 |
| 276 | L | 0.96460177 | I | 0.00884956 | ND | 0.02654867 |
| 278 | N | 0.04424779 | ND | 0.02654867 | A | 0.92920354 |
| 303 | V | 0.00884956 | I | 0.92035398 | ND | 0.07079646 |
| 304 | L | 0.01769912 | I | 0.91150442 | ND | 0.07079646 |
| 317 | V | 0.92477876 | I | 0.00442478 | ND | 0.07079646 |
| 326 | N | 0.00442478 | ND | 0.07079646 | T | 0.92477876 |
| 342 | N | 0.07964602 | F | 0.84955752 | ND | 0.07079646 |
| 363 | K | 0.0619469 | S | 0.78761062 | ND | 0.15044248 |
| 367 | L | 0.03539823 | D | 0.75221239 | ND | 0.21238938 |
| 368 | D | 0.03097345 | ND | 0.32300885 | T | 0.6460177 |
| 369 | ND | 0.3539823 | L | 0.05309735 | P | 0.42477876 |
| 426 | L | 0.00442478 | I | 0.48230088 | ND | 0.51327434 |

**Table 3:** Segregating sites in Pv47 globally. Amino acids in bold are those found in samples in this study. ND represents no data.

| **Position** | **Amino Acid** | **Frequency** | **Amino Acid** | **Frequency** | **Amino Acid** | **Frequency** | **Amino Acid** | **Frequency** |
| --- | --- | --- | --- | --- | --- | --- | --- | --- |
| **22** | **L** | **0.95588** | F | 0.04412 |  |  |  |  |
| **24** | **L** | **0.96324** | F | 0.03676 |  |  |  |  |
| **27** | **E** | **0.96324** | K | 0.03676 |  |  |  |  |
| 29 | L | 0.99265 | I | 0.00735 |  |  |  |  |
| 31 | N | 0.00735 | D | 0.99265 |  |  |  |  |
| 57 | T | 0.40441 | S | 0.59559 |  |  |  |  |
| 62 | S | 0.97794 | N | 0.02206 |  |  |  |  |
| 66 | Y | 0.99265 | H | 0.00735 |  |  |  |  |
| 80 | T | 0.99265 | P | 0.00735 |  |  |  |  |
| 82 | L | 0.97794 | V | 0.02206 |  |  |  |  |
| 98 | N | 0.00735 | Y | 0.99265 |  |  |  |  |
| 156 | D | 0.91912 | G | 0.02206 | ND | 0.05882 |  |  |
| 193 | T | 0.93382 | I | 0.00735 | ND | 0.05882 |  |  |
| 230 | I | 0.11765 | ND | 0.05882 | V | 0.82353 |  |  |
| 233 | M | 0.36765 | I | 0.57353 | ND | 0.05882 |  |  |
| 237 | F | 0.91912 | I | 0.02206 | ND | 0.05882 |  |  |
| 240 | E | 0.88971 | D | 0.05147 | ND | 0.05882 |  |  |
| 262 | T | 0.10294 | I | 0.12500 | K | 0.71324 | ND | 0.05882 |
| 273 | M | 0.00735 | I | 0.88971 | ND | 0.05882 | V | 0.04412 |
| 337 | F | 0.00735 | ND | 0.05882 | C | 0.93382 |  |  |
| 373 | A | 0.71324 | ND | 0.05882 | V | 0.22794 |  |  |

**Table 4:** Presence of homozygous allele and allele frequency of histidine and glutamine at amino acid position 53 in P47 receptor over time.

|  | **Semera 2018** | **Kebridehar 2018** | **Dire Dawa 2018** | **Lawyacado 2021** | **Dire Dawa 2022** | **Dire Dawa 2022 Malaria Positive** | **India 2022** |
| --- | --- | --- | --- | --- | --- | --- | --- |
| **HH** | 14 | 14 | 14 | 8 | 13 | 6 | 12 |
| **QQ** | 0 | 0 | 2 | 1 | 1 | 0 | 1 |
| **HQ** | 3 | 2 | 7 | 2 | 5 | 0 | 7 |
| **Total:** | 17 | 16 | 23 | 11 | 19 | 6 | 20 |
| **Frequency of Q** | 0.088235 | 0.0625 | 0.23913 | 0.181818182 | 0.184211 | 0 | 0.225 |
| **Frequency of H** | 0.911765 | 0.9375 | 0.76087 | 0.818181818 | 0.815789 | 1 | 0.775 |
